# Size conservation emerges spontaneously in biomolecular condensates formed by scaffolds and surfactant clients

**DOI:** 10.1101/2021.04.30.442154

**Authors:** Ignacio Sanchez-Burgos, Jerelle A. Joseph, Rosana Collepardo-Guevara, Jorge R. Espinosa

## Abstract

Biomolecular condensates are liquid-like membraneless compartments that contribute to the spatiotemporal organization of proteins, RNA, and other biomolecules inside cells. Some membraneless compartments, such as nucleoli, are dispersed as different condensates that do not grow beyond a certain size, or do not present coalescence over time. In this work, using a minimal protein model, we show that phase separation of binary mixtures of scaffolds and low-valency clients that can act as surfactants—i.e., that significantly reduce the droplet surface tension—can yield either a single drop or multiple droplets that conserve their sizes on long timescales (herein ‘multidroplet size-conserved’), depending on the scaffold to client ratio. Our simulations demonstrate that protein connectivity and condensate surface tension regulate the balance between these two scenarios. Multidroplet size-conserved behavior spontaneously arises at increasing surfactant-to-scaffold concentrations, when the interfacial penalty for creating small liquid droplets is sufficiently reduced by the surfactant proteins that are preferentially located at the interface. In contrast, low surfactant-to-scaffold concentrations enable continuous growth and fusion of droplets without restrictions. Overall, our work proposes one potential thermodynamic mechanism to help rationalize how size-conserved coexisting condensates can persist inside cells—shedding light on the roles of general biomolecular features such as protein connectivity, binding affinity, and droplet composition in this process.

## Introduction

To fulfill their biological functions, cells must organize their contents into different compartments. One way of achieving spatiotemporal organization is via the formation of membrane-bound organelles such as the nucleus^1^ or lysosomes^2^; while another is by self-assembly of proteins and nucleic acids through a process called liquid–liquid phase separation (LLPS)^3–6^, as in stress granules^7^, P granules^8^ or nuclear speckles^9^. Significant interest in biomolecular LLPS has recently emerged due to wide-ranging implications on cellular function, including genome silencing^10,11^, signaling^12^, buffering cellular noise^13^, and formation of super-enhancers^14^, among many others^15,16^. Furthermore, neurodegenerative diseases such as Alzheimer’s, Parkinson’s or ALS have been also associated with aberrant liquid-to-solid transitions from phase-separated compartments^17–19^. In LLPS, protein-rich liquid condensates coexist with a protein-poor liquid matrix. The chief drivers of biomolecular LLPS are multivalent proteins (i.e., those possessing multiple interaction sites)—widely known as ‘scaffolds’^20^. Scaffold proteins typically show high affinity for other biomolecules, including ‘client’ molecules. Unlike scaffolds, clients are unable to undergo LLPS on their own and are instead recruited to the condensates by interacting with scaffolds. Together, scaffolds and clients shape the biomolecular condensates that play fundamental roles within the cell^21–24^.

A crucial open question regarding the emergence of phase-separated condensates is the existence of compartments that do not grow beyond a given size^25–28^. Basic thermodynamics suggest that over time, LLPS should result in the formation of a single large condensate rather than multiple coexisting small droplets^29^. The latter case is disfavored because it yields an overall high droplet surface area to volume ratio (i.e., high interfacial penalty), while the former ensures that the interfacial free energy penalty is minimized. Despite of the thermodynamic preference for single condensates over multidroplet systems, both types of architectures can be present within cells. Indeed, single large condensates have been observed^22,30^, but also diverse coexisting size-restricted droplets have been reported in different *in vivo* systems, such as in the amphibian oocyte nucleolus^27^, in the *Saccharomyces cerevisiae* cytoplasmic processing bodies^31^ and elsewhere^32–34^. Additionally, size-conserved multidroplet architectures have been found in non-coalescing ribonucleoprotein condensates^22^ and in multiphase complex coacervates *in vitro^35^.* However, the underlying molecular mechanisms and driving forces behind the formation of multiple coexisting droplets, also known as emulsification^36^, require further investigation.

Several explanations for why or how emulsions can be thermodynamically stable at biological relevant time-scales are currently under debate^37^. One possible mechanism is that the presence of active ATP-dependent processes might conveniently regulate the conditions where droplets grow and coalesce^38–40^. Other studies suggest that proteins with various highly distinct interacting domains may form micelle-like condensates^28,41,42^. In multicomponent mixtures, another possibility could be that specific binding proteins act as powerful surfactants, and thus, modulate the droplet surface tension penalty leading to multicondensate coexistence^32,33,35^. Moreover, a recent alternative explanation suggests that the interplay between protein diffusion and saturation of protein binding sites can also induce size-conserved condensate formation^25,26^. It is plausible that all these different mechanisms contribute to the size conservation of condensates inside cells under different conditions. In this work, we use a minimal protein model, which recapitulates the relationship between protein valency and critical parameters^43–45^, to investigate the regulation of droplet size in binary mixtures of multivalent proteins (scaffolds and clients). We show that liquid–liquid phase separation of scaffolds and clients mixtures, where clients act as surfactants, can give rise to condensates forming single droplets or multiple size-conserved droplets (Fig. 1 (a)), and that the transition between the two scenarios can be regulated by the condensate scaffold/surfactant client ratio. Furthermore, our simulations suggest how general molecular features such as protein connectivity, binding affinity, and droplet composition can critically modulate and stabilize the formation of size-conserved condensates^25,26^, and might have also implications to understanding the phase behavior of multilayered condensates^35,45,46^.

**Figure 1.**
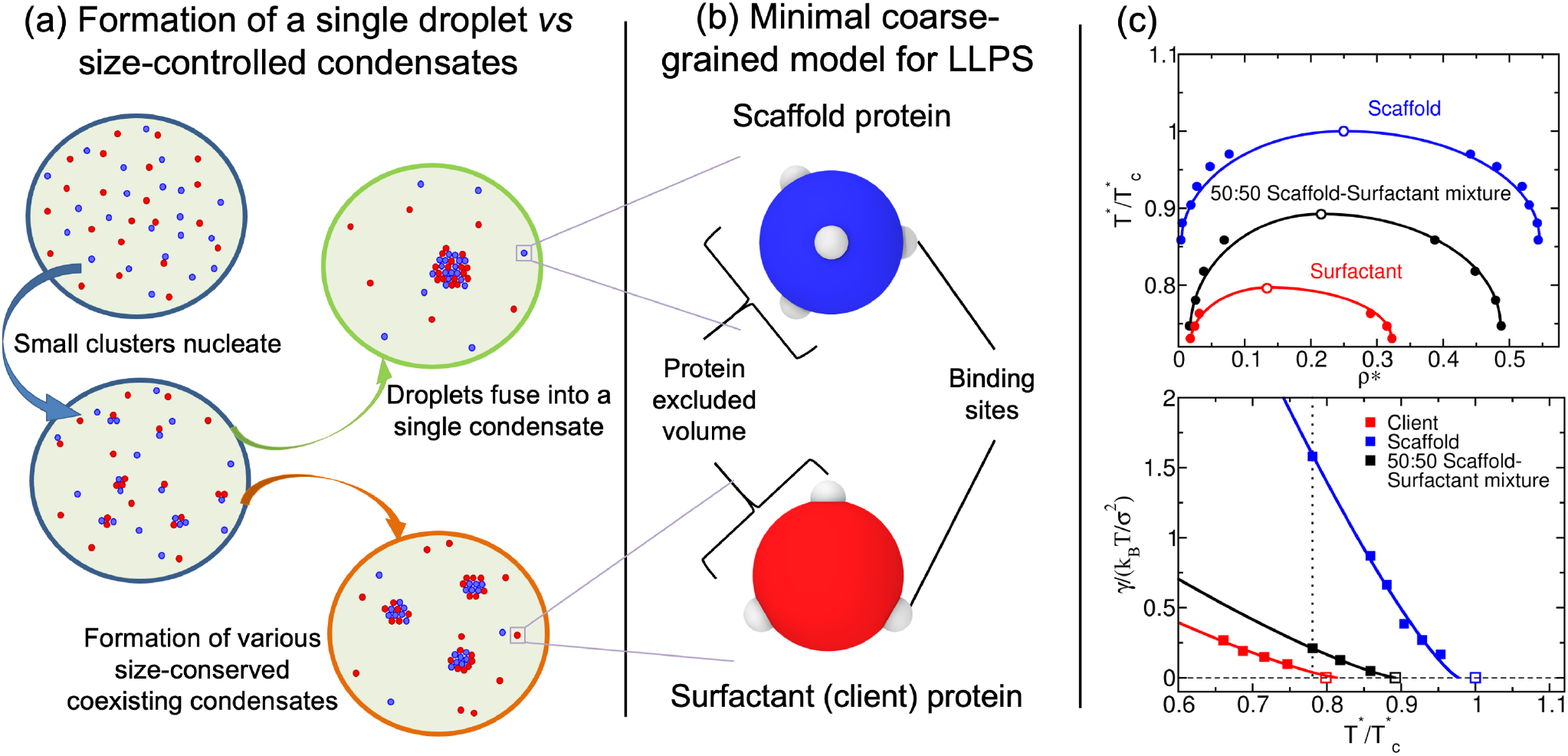
(a) Schematic representation of single-droplet formation versus size-conserved protein condensation. Scaffold proteins are depicted as blue spheres, while surfactant clients are shown as red spheres. (b) Minimal coarse-grained model for protein LLPS: Blue and red spheres represent the excluded volume of scaffold and surfactant (client) proteins respectively, while gray patches represent the binding sites of the proteins. Two different proteins are modeled: scaffold proteins, with 4 promiscuous binding sites in a tetrahedral arrangement, and surfactant proteins, with 3 binding sites in a planar equidistant arrangement that can only bind to scaffold binding sites. Details on the model parameters are provided in the Methods section. (c) Top: Phase diagram in the temperature–density plane for a scaffold protein system (blue), a 50:50 scaffold–surfactant mixture (black) and for a hypothetical surfactant system in which client proteins can self interact (red). The latter system further illustrates the effect of multivalency in LLPS. Filled circles indicate the estimated coexisting densities from Direct Coexistence simulations^66–68^, and empty ones depict the critical points calculated via the law of rectilinear diameters and critical exponents^69^ (see Supporting Information for details on these calculations). Bottom: Surface tension (*γ*) dependence on temperature for the scaffold system (blue), the 50:50 scaffold–surfactant mixture (black), and the hypothetical system in which surfactant proteins can self interact (red). Filled squares represent direct estimations of *γ*, continuous curves depict fits to our data of the following form: 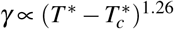, and empty squares show the critical temperature of each system evaluated through the law of rectilinear diameters and critical exponents^69^. The vertical dashed line indicates the temperature at which the remainder of our study is performed. Note that temperature (in reduced units, *T*\*) is renormalized by the critical temperature 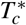 (also in reduced units) of the scaffold protein system 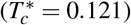.

## Results

### A minimal protein model for scaffold and client mixtures

Coarse-grained (CG) potentials have emerged as powerful tools for describing the phase behavior of biomolecules, such as proteins and nucleic acids, and delineating the underlying physicochemical features that drive LLPS^47,48^. Various levels of molecular resolution can be achieved with CG models; encompassing mean field models^49,50^, lattice-based simulations^51,52^, minimal models^44,45,53–56^, and coarse-grained sequence-dependent simulations^42,57–61^. Here, we employ our minimal protein model^43^, which has been previously applied to unveil the role of protein multivalency in multicomponent condensates^62^ and multilayered condensate organization^63^, as well as to investigate the role of RNA in RNA-binding protein nucleation and stability^64^. In this model, proteins are described by a pseudo hard-sphere potential^65^ that accounts for their excluded volume, and by short-range potentials for modeling the different protein binding sites, and thereby mimicking protein multivalency^43^ (Fig. 1 (b)). For computational efficiency, an implicit solvent is used, and therefore, the condensed phase corresponds to a liquid phase, while the protein diluted phase is essentially a vapor phase. In what follows, the unit of distance is σ, the molecular diameter of the proteins (both scaffolds and clients), and the unit of energy *k_B_T* (for further details on the model parameters and the employed reduced units see the Methods section).

We define scaffolds as proteins that can establish both homotypic interactions and heterotypic interactions with clients, while clients (or hereafter called surfactants) as proteins that are limited to bind only to scaffolds and do not bind to other clients (following the original definition of Banani *et al*.^20^). Within this scheme, phase separation is driven by scaffolds (high-valency proteins), whereas surfactants (proteins with lower valency) are recruited to condensates at the expense of depleting LLPS-stabilizing scaffold–scaffold interactions. The two-phase coexisting densities as a function of temperature for our model of scaffold proteins (blue), a 50:50 binary mixture of scaffolds and surfactants (black), and a system composed just by surfactants (red) are depicted in Fig. 1(c) (Top panel). Note that in the 50:50 mixture, surfactant proteins do not homotypically interact, while in the pure surfactant system – for comparison purposes on the effect of protein valency – surfactant proteins can self-interact. The addition of surfactant proteins that are strong competitors for the scaffold binding sites significantly hinders the ability of scaffolds to phase separate (i.e., clients lower the critical temperature)^20,62^. Moreover, the presence of surfactant clients drastically reduces the surface tension of the condensates (black curve) as shown in Fig. 1(c) (Bottom panel). In the following section, we elucidate the implications of the client-induced surface tension reduction on the behavior of phase-separated condensates.

### Size-conserved condensates can be regulated by surfactant proteins

Despite their seemingly subordinate role in condensate formation, client molecules (e.g. low valency proteins or self-avoiding nucleic acids) can significantly impact the organization of biomolecular condensates^20,35,70,71^. Remarkable examples include the multilayered organization of stress granules^72,73^, the nucleoli^22^, nuclear speckles^74^, mixtures of RNA-binding proteins with RNA^75,76^, and *in vitro* complex coacervates^35,70,71^. While scaffolds mainly contribute to maximize the condensate liquid network connectivity^20,62^, clients can severely reduce the droplet interfacial free energy penalty of creating an interface^45^. The proliferation marker protein Ki-67, a component of the mitotic chromosome periphery, is an important example of a natural surfactant in intracellular compartmentalization^32^. Our simulations also show that the recruitment of clients to the condensate, that can act as surfactants, greatly decreases its surface tension (Fig. 1(c) (Bottom)).

To further investigate the role of surfactant clients, we perform a set of Direct Coexistence simulations^66–68^ for the pure scaffold system and the 50:50 binary mixture of scaffold and surfactants shown in Fig. 1(c). For a constant temperature of 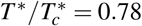 and a global (system) density of *ρ*\*=0.136 (both in reduced units; see Methods section for further details on the employed reduced magnitudes), we create three simulation box geometries with the dimensions summarized in Table 1. Using this approach, we can effectively modulate the surface/volume ratio of the condensed phase (hereafter called droplet): where the droplet surface is *S* = 2 * *L_y_* * *L_z_* and its volume is *V = L_y_* * *L_z_ * L_x,slab_*; with *L_x,slab_* representing the width of the condensate in the *x* direction. Fig. 2(a)–(c), summarizes the phase behavior of the pure scaffold system (Top panels) and the 50:50 binary mixture (Middle panels) along the three designed simulation box geometries. The pure scaffold condensate exhibits distinct surface/volume ratios depending on the box geometry (see Table 1), having all of them the same droplet density. In other words, the scaffold condensate would be able to continuously grow as a single droplet at expense of the diluted phase until reaching equilibrium. In contrast, the scaffold-surfactant mixture yields various coexisting equilibrium droplets with a roughly constant surface/volume ratio of 0.21(2) σ^-1^ in all systems (Table 1). This result strongly suggests how size-conserved multidroplet phase behavior can be simply induced by the presence of low valency clients, which actively contribute to lowering the surface tension of the droplets (Fig. 1(c) (Bottom)) - thereby serving as natural surfactants^32,33^.

**Figure 2.**
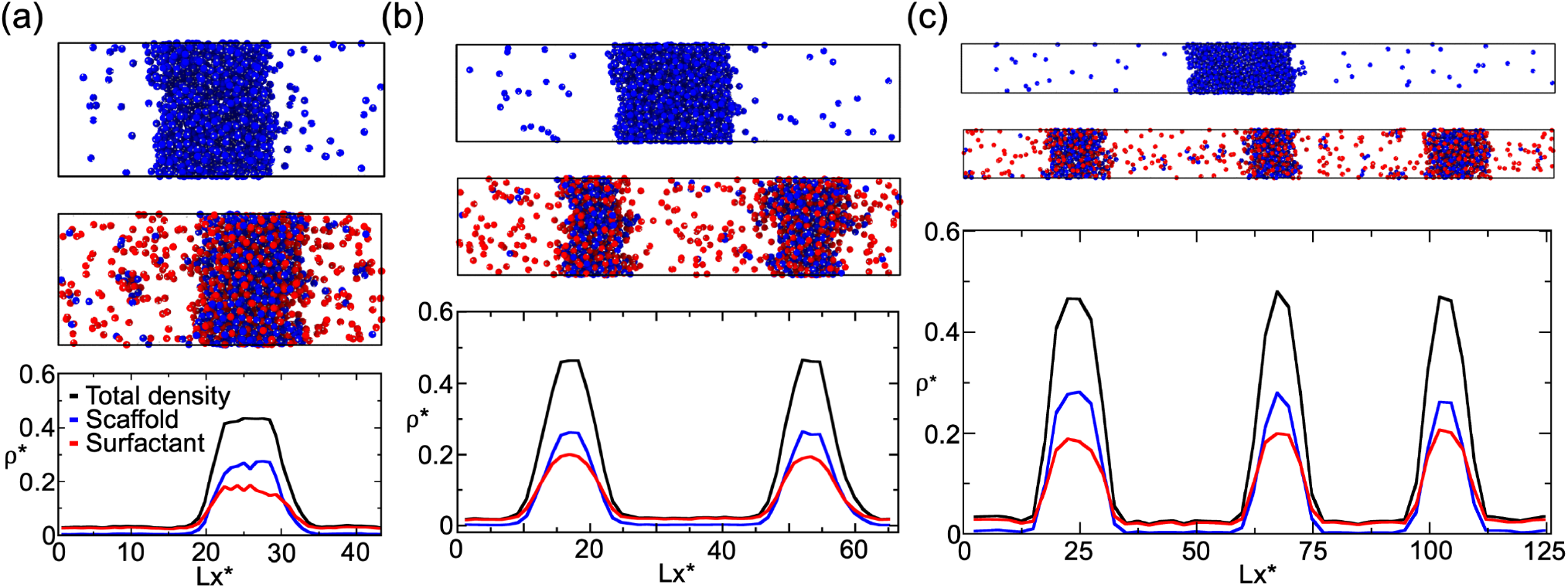
Direct Coexistence simulations for a scaffold protein system (Top panels) and a 50:50 binary mixture of scaffold and surfactant proteins (Middle panels) with different simulation box geometries (see Table 1 for the different simulation box sides employed in *a, b* and *c* geometries). All simulations were performed at 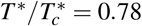 and a global density of *ρ*\* =0.136. (a)–(c) (Top and Middle panels): Scaffold proteins are colored in blue and surfactant clients in red; the protein binding sites of both protein types are colored in gray. (a)–(c) (Bottom panels): Density profiles along the long side (x) of each simulation box for the 50:50 scaffold–surfactant mixture. The (scaffold+surfactant) total density profile is depicted in black, the surfactant client density profile is shown in red, and that of scaffolds in blue. Densities of the different scaffold–surfactant coexisting droplets as well as the scaffold molar fraction in the condensed and diluted phase for each geometry are given in Table 2. The total density of the pure scaffold condensates in all geometries is 0.54(1).

**Table 1.** Simulation box dimensions and condensate surface/volume ratios of the three systems represented in Fig. 2. Geometries (a), (b) and (c) account for the (a), (b) and (c) panels shown in Fig. 2 respectively.

| Box geometry | (a) | (b) | (c) |
| --- | --- | --- | --- |
| $L_x / \sigma$ | 43.7 | 68.1 | 124.8 |
| $L_y=L_z / \sigma$ | 17.5 | 13.7 | 10.3 |
| Scaffold S/V Ratio / $\sigma^{-1}$ | 0.14(2) | 0.12(2) | 0.09(2) |
| 50:50 Mixture S/V Ratio / $\sigma^{-1}$ | 0.21(2) | 0.21(2) | 0.20(2) |

We analyze the composition of the different coexisting scaffold-surfactant condensates along the distinct box geometries (Fig. 2 (Bottom panels)). In all cases, the properties of the droplets are remarkably similar; this is also the case for the pure scaffold condensates. The density of all droplets, as well as their condensate composition and surface tension is roughly constant (Table 2). Notably, droplet density profiles reveal how the surfactant profile decays considerably slower than that of the scaffold proteins from the core condensate; indicating that the surfactant partition coefficient at the interface is greater than that of the scaffolds. Previous works suggest that this behavior is preferable as it minimizes the condensate surface tension^32,35,63^. We also verify that the coexisting droplets are thermodynamically stable states, rather than just metastable, by simulating long timescales to allow for droplet fusion or variation of their composition. However, even at a close distance, droplets coexist without coalescing or altering their equilibrium composition. The multidroplet behavior of size-conserved condensates (in our case of ~0.21 σ^-1^ surface/volume ratio) is a consequence of the thermodynamic conditions of our system (i.e., mixture composition, temperature, and density). Note that droplet curvature effects such as Laplace internal pressure^77^ or surface tension dependence on droplet curvature^78^ have not been considered in our simulations, since we do not expect them to play an important role at biologically relevant droplet size scales (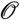 *μ*m)^37^. Those effects are only dominant in the nanometer scale (i.e., up to droplet radii of tens of nanometers)^79–81^.

**Table 2.** Properties of systems presented in Fig. 2 containing 50:50 scaffold-surfactant compositions in different simulation box geometries. In all cases, 6 independent simulations (with different initial velocity distributions), each starting from a pre-equilibrated configuration, were performed. For the geometries with more than one droplet, the values are averaged over the different droplets, although the variance between distinct droplets is significantly small as shown in Fig. 2 (Bottom panels).

| Box geometry | (a), 1 droplet | (b), 2 droplets | (c), 3 droplets |
| --- | --- | --- | --- |
| $\gamma / (k_B T / \sigma^2)$ | 0.26(4) | 0.23(4) | 0.23(4) |
| Scaffold molar fraction in condensate | 0.585(5) | 0.576(5) | 0.583(5) |
| Scaffold molar fraction in diluted phase | 0.041(2) | 0.041(2) | 0.043(2) |
| $\rho_{\text{condensate}}^*$ | 0.47(1) | 0.47(1) | 0.46(1) |
| $\rho_{\text{diluted phase}}^*$ | 0.030(3) | 0.027(3) | 0.029(3) |

The presence of surfactant clients within the condensate substantially lowers the liquid network connectivity^62,82,83^, and therefore, reduces the enthalpic gain sustaining LLPS. Consequently, the system minimizes its free energy by optimizing the number of surfactants that are incorporated into the bulk condensed phase; i.e., by creating higher surface/volume ratios, where surfactants are preferentially located towards the interface rather than in the core. Such free energy optimization yields multiple coexisting condensates of a certain size, rather than a single-condensate system. This behavior is only thermodynamically favorable when the condensate surface tension is very low, as in the case of 50:50 scaffold-surfactant condensates with a surface tension of *γ* = 0.23 *k_B_T σ*^-2^—almost 7 times lower than that of the pure scaffold condensate (*γ* = 1.58 *k_B_T σ*^-2^) at the same conditions 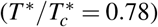.

### Surfactant concentration critically modulates droplet size

As discussed in the previous section, our minimal protein model shows that for a given composition of scaffold and surfactant clients, independently of the imposed box geometry, droplets can only grow to a certain size. This size-restricted growth is, in turn, determined by the optimal surface/volume ratio that minimizes the free energy of the two coexisting phases. Therefore, larger system sizes lead to higher number of coexisting size-conserved droplets. On the other hand, when the condensate is only composed of scaffold proteins, as the system size increases, the size of the condensate simply grows instead of yielding new multiple size-conserved droplets (Fig. 2). These results illustrate how a simple model for scaffold and surfactant proteins, merely controlled by protein valency and binding affinity, can recapitulate mesoscale features of *in vivo* and *in vitro* condensates that exhibit size-conserved growth^22,27,31–35^. Since this phase behavior only arises when the concentration of surfactants is not negligible, we now investigate how condensates can switch between both scenarios, and how their surface/volume ratio is modulated by their relative scaffold-surfactant composition.

By gradually increasing the surfactant client concentration of the scaffold–surfactant mixture (at a constant temperature and system density), the condensate size progressively decreases to accommodate the equilibrium droplet surface/volume ratio at those conditions to the simulation box geometry; i.e., the condensate splits into two, and subsequently into three coexisting liquid droplets (see Fig. 3). We note that the composition of the different coexisting droplets at a given surfactant concentration is remarkably similar, highlighting that all droplets are in equilibrium. In parallel, we evaluate the surface tension of the different coexisting droplets as a function of surfactant concentration. We find that *γ* monotonically decreases (but not linearly) as the surfactant concentration increases (Fig. 3). This is not surprising (see Fig. 1 c) given that one of the key molecular driving forces behind size-conserved multidroplet formation is the reduction of *γ* by surfactants coating the droplet surface. Above a certain surfactant concentration – > 65% for our system and given conditions – LLPS is inhibited. Beyond this limit, the condensate liquid network connectivity sustained by scaffold proteins can no longer compensate the mixing entropy of the system. We also analyze the surface/volume ratio of the droplets as a function of surfactant concentration. At infinitely low surfactant concentration, the condensate displays a ratio of M).09 σ^-1^; however, this ratio is fully determined by the total number of proteins in the system and the box geometry, since as shown in Fig. 2, scaffold condensates can reach any droplet size when surfactants are absent. At low surfactant concentrations (i.e., % surfactant < 27.5%), the maximum droplet size corresponds to a surface/volume ratio of ~0.11 σ^-1^. Beyond that threshold concentration, the condensate slab shrinks, and to achieve their equilibrium surface/volume ratio, it splits into smaller coexisting droplets. The maximum equilibrium droplet size in the two-droplet regime is that corresponding to ratios of ~0.19 σ^-1^. Finally, for surfactant compositions higher than 38%, three coexisting condensates emerge. The maximum surface/volume ratio that droplets can achieve is ~0.27 σ^-1^ at 60% client composition, which is only possible due to the extreme reduction in the surface tension (*γ* = 0.15 *k_B_T σ*^-2^) - more than one order of magnitude lower than that of the pure scaffold condensate (γ = 1.58 *k_B_T σ*^-2^) at the same temperature and system density.

**Figure 3.**
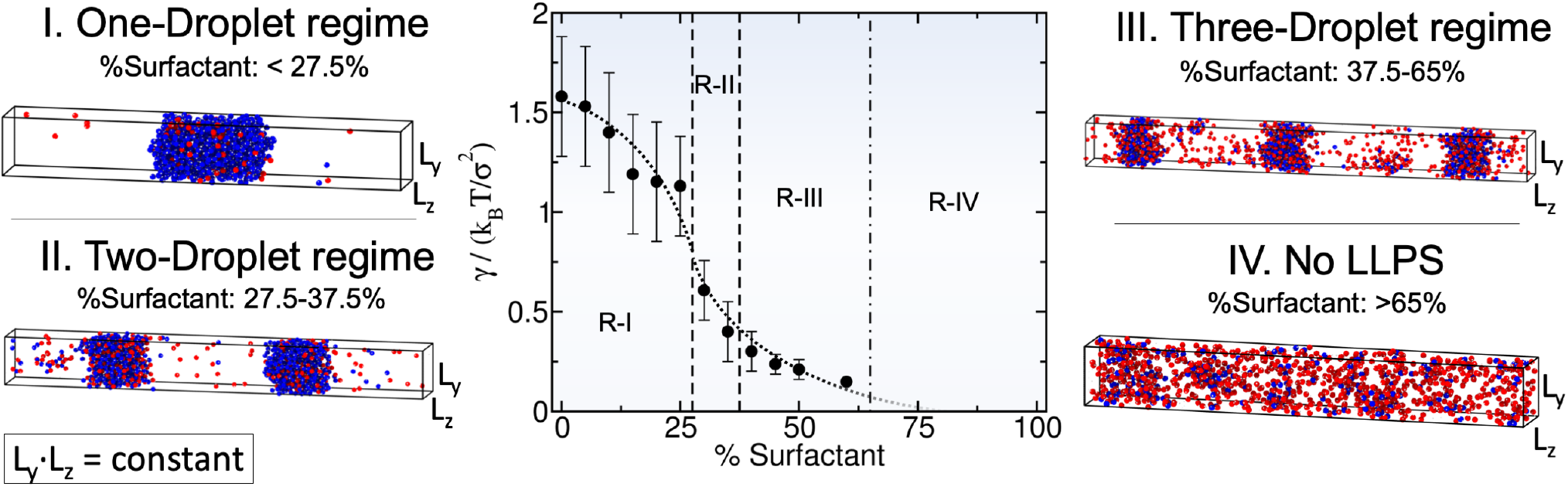
Surface tension (*γ*) dependence on the scaffold-surfactant composition (in %) at 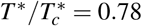 and system (global) density of *ρ*\* =0.136. Vertical dashed lines indicate the maximum surfactant concentration which allows LLPS for a given number-droplet regime in our system. Note that the maximum droplet size varies continuously with surfactant concentration even within the same number-droplet regime. The maximum condensate size in terms of surface/volume ratio along the different number-droplet regimes are: ~0.11 σ^-1^ for the one-droplet regime, ~0.19 σ^-1^ for the two-droplet regime, and ~0.27 σ^-1^ for the three-droplet regime. At surfactant client compositions exceeding 65% there is no LLPS. For each computed composition, the global density of the system and the simulation box cross-section *(L_z_L_y_*) are kept constant. Snapshots of the DC simulations along different surfactant composition droplet regimes are included. Please note that in the multidroplet regime, the different coexisting droplets exhibit similar compositions and densities.

Previous studies have highlighted the challenges associated with measuring condensate surface tensions due to the small size of protein droplets^6,84^. Nonetheless, there are available estimates of this magnitude for ribonucleoprotein condensates, and these measurements demonstrate that surfactant proteins can reduce *γ* by orders of magnitude^22^. With our minimal model, we qualitatively observe such behavior when surfactant proteins are recruited to the condensate, giving rise to emulsions of multiple coexisting droplets with very low surface tension. Surfactant proteins can lead to the formation of multidroplet emulsions by inducing multilayered condensate architectures^22,35^. Diverse biomolecular organelles such as stress granules^72,73^, the nucleoli^22^, or nuclear speckles^74^ exhibit this organization. Moreover, different *in vitro* complex coacervates^35,46,70,71^, and mixtures of RNA-binding proteins and RNA molecules^75,76^ are also known to show multilayered assembling. In Fig. 4(a), we analyze the droplet architecture of a protein condensate with a 50:50 scaffold-surfactant composition (i.e., the system in Fig. 2(a) (Middle panel)). We find that in the droplet core, the scaffold to surfactant relative abundance is much higher than along the interfacial region (Fig. 4 a), where it drops to almost half than that of the surfactant proteins. Nonetheless, the surfactant concentration within the droplet core is still remarkably high considering its destabilizing role in the condensate liquid network connectivity^44^. The observed non-homogeneous condensate organization stems from the higher valency of scaffold proteins, which allows them to establish higher molecular connectivity within the core condensate, and thus, induce higher enthalpic gain upon multilayered assembly.

**Figure 4.**
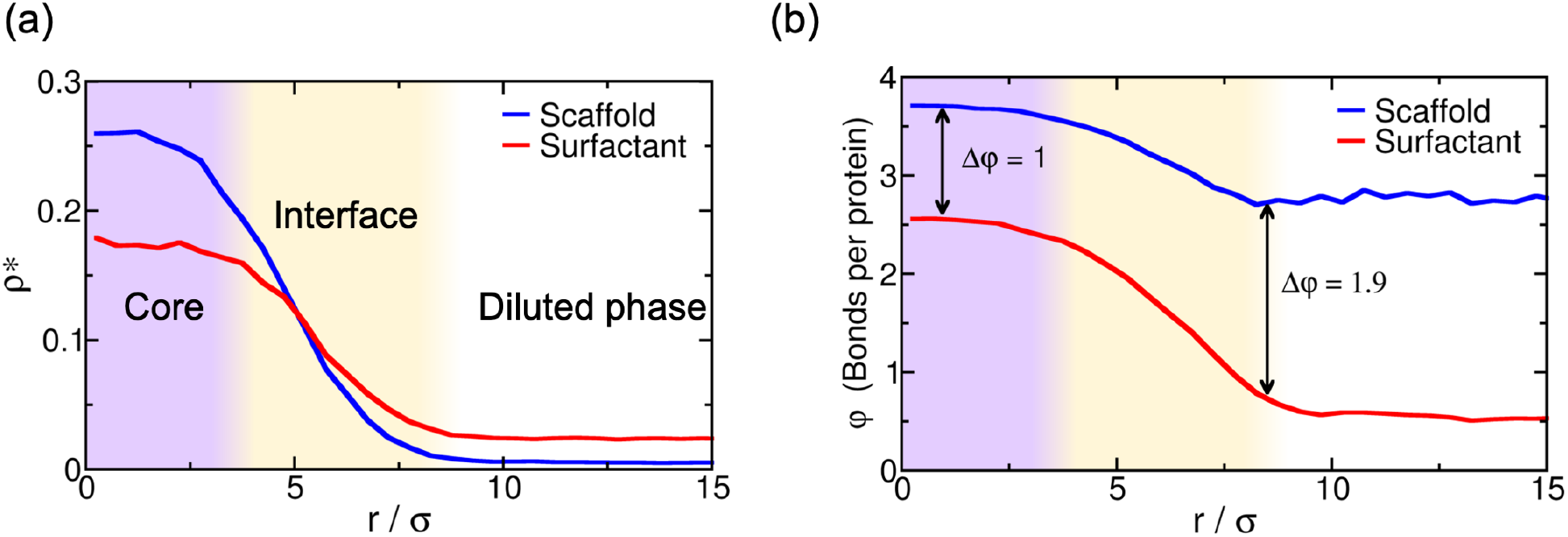
(a) Droplet density profile (in reduced units) for scaffold (blue) and surfactant proteins (red) from the droplet center of mass towards the surrounding diluted phase for the 50:50 binary mixture within the simulation box geometry shown in Fig. 2(a) at 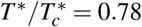. (b) Average number of engaged binding sites per protein (*φ*) as a function of distance from the droplet center of mass for scaffold (blue) and surfactant proteins (red). One binding site is considered to be engaged to another if the distance between them is less than 0.145 σ (i.e., the maximum bond length interaction between distinct protein binding sites; for further details on these calculations see Supporting Information).

Finally, to gain further insight on the droplet liquid network connectivity, we evaluate the average number of engaged binding sites per protein (*φ*) as a function of distance from the center of mass of the condensates (Fig. 4(b)). Scaffold proteins present a significantly higher amount of molecular connections per protein than surfactants (i.e., *φ* ~ 3.7 and *φ* ~ 2.5 for scaffolds and surfactants, respectively), at the droplet core. This observation highlights how surfactants negatively contribute to the stability of the condensate. However at the interface, such diminished connectivity of surfactant proteins (*φ* ~ 1) with respect to that of scaffolds (*φ* ~ 3) substantially reduces the condensate surface tension - by decreasing the enthalpic cost (Δ*h_i_*) of creating an interface. This energetically favorable protein arrangement, controlled in our model just by the variance in protein valency of the components, is expected to be contributed also by changes in relative binding affinities among proteins, and be modulated by post-translational modifications *in vivo*^21,31^. Furthermore, such variations can be also relevant to understand the physical parameters controlling multilayered condensate organization and, ultimately, regulation of the formation of size-conserved multidroplet emulsions^25,26,28^.

## Conclusions

In this work, we employ our minimal protein model^43^ to demonstrate how biomolecular multidroplet emulsification can be controlled by the subtle balance between liquid network connectivity and droplet surface tension, and how general molecular features such as protein valency and binding affinity can critically regulate this behavior. By using a binary mixture of scaffold and client proteins that act as surfactants (following the original definition proposed by Banani *et al*.^20^), we design a set of Direct Coexistence simulations in which we can conveniently modulate the simulation box geometry to assess the propensity of the condensates to accommodate different surface/volume ratios. The ability (or disability) of these mixtures to adopt different surface/volume ratios, imposed by the box geometry, can be regarded as an indirect measurement of the droplet propensity to grow beyond a certain size. We find that pure component scaffold condensates can easily adapt to distinct surface/volume ratios; in support of their ability to grow and fuse into a single droplet. However, 50:50 binary scaffold–surfactant mixtures stabilize instead several coexisting liquid condensates with roughly constant surface/volume ratios to accommodate the imposed system size geometry. Such behavior is a clear signature of size-conserved multidroplet emulsification, as found in the nucleolus^27^, ribonucleoprotein condensates^22^, micelle-like condensates^28,41^, and *in vitro* complex coacervates^35^.

We also elucidate the role of surfactant concentration in size-conserved droplet growth. By gradually decreasing the scaffold–surfactant ratio in our mixtures, we observe that the maximum droplet size is reduced, while simultaneously increasing the number of coexisting condensates. This trend continues until a sufficiently high surfactant concentration is reached, where LLPS is no longer possible. Moreover, as clients are added, the droplet surface tension dramatically decreases, facilitating the formation of multiple coexisting small liquid droplets at low interfacial energetic cost. Client proteins, besides decreasing the stability of the condensates^62^, can effectively behave as natural droplet surfactants^32,33^. Due to their considerably lower molecular connectivity compared to that of scaffolds, surfactants preferentially migrate towards the droplet interface; thereby, minimizing the enthalpic cost of creating an interface^45^. Heterogeneous molecular organizations of condensates have been observed in stress granules^72,73^, the nucleoli^22^ and nuclear speckles^74^. We find that such heterogeneity enable the maximization of the condensate liquid network connectivity through scaffold–scaffold protein interactions within the droplet core.

Rationalizing the underlying mechanisms employed by cells to precisely regulate the size of their diverse membraneless compartments and processing bodies^27,31,33^ represents a crucial step towards understanding intracellular spatiotemporal cell organization. Taken together, our coarse-grained simulations help to elucidate the relationship between single-droplet phase formation and size-conserved multidroplet architecture, and put forward general molecular features such as valency and binding affinity as chief drivers in these scenarios.

## Methods

We model our coarse-grained multivalent proteins using the MD-Patchy potential proposed in Ref.^43^, which is composed by two different set of potentials: a Pseudo Hard-Sphere (PHS) potential^85^ to continuously describe the repulsive interaction and excluded volume between different protein replicas, and a continuous square-well (CSW)^86^ potential to describe the patch-patch interactions among different protein binding sites. The *uPHS* potential is described by the following expression:

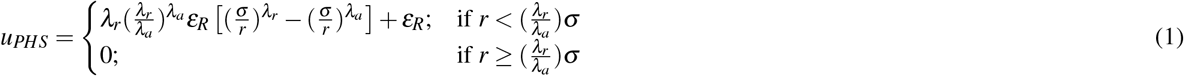

where *λ_a_* = 49 and *λ_r_* = 50 are the exponents of the attractive and repulsive terms respectively, *ε_R_* accounts for the energy shift of the PHS interaction, *σ* is the molecular diameter (and our unit of length) and *r* is the center-to-center distance between different PHS particles. For the patch-patch interaction we use the following expression:

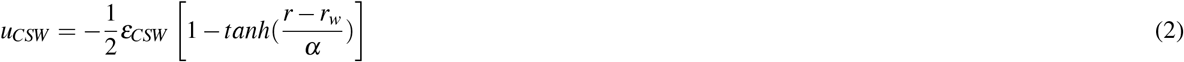

where *ε_CSW_* is the depth of the potential energy well, *r_w_* the radius of the attractive well, and *α* controls the steepness of the well. We choose *α* = 0.005σ and *r_w_* = 0.12σ so that each patch can only interact with another single patch.

The mass of each patch is a 5% of the central PHS particle mass, which is set to 3.32×10^-26^kg, despite being this choice irrelevant for equilibrium simulations. This 5% ratio fixes the moment of inertia of the patchy particles (our minimal proteins). The molecular diameter of the proteins, both scaffold and clients, is σ=0.3405 nm, and the value of *ε_R_/k_B_* is 119.81K. All our results are presented in reduced units: reduced temperature is defined as *T*\* = *k_B_T/ε_CSW_*, reduced density as *ρ*\* = (*N/V*)*σ*^3^, reduced pressure as *p*\* = *pσ*^3^/(*k_B_T*), and reduced time as 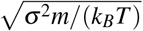. In order to keep the PHS interaction as similar as possible to a pure HS interaction, we fix *k_B_T/ε_R_* at a value of 1.5 as suggested in Ref.^85^ (fixing *T* = 179.71K). We then control the effective strength of the binding protein attraction by varying *ε_CSW_* such that the reduced temperature, *T*\* = *k_B_T/ε_CSW_*, is of the order of 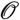(0.1).

Since both *u_PHS_* and *u_CSW_* potentials are continuous and differentiable, we perform all our simulations using the LAMMPS Molecular Dynamics package^87^. Periodic boundary conditions are used in the three directions of space. The timestep chosen for the Verlet integration of the equations of motion is Δ*t*\*=3.7×10^-4^. The cut-off radius of the interactions of both potentials is set to 1.17σ. We use a Nosé-Hoover thermostat^88,89^ for the *NVT* simulations with a relaxation time of 0.074 in reduced units. For *NpT* simulations, a Nosé-Hoover barostat is employed with the same relaxation time^90^.

The methodological details of the calculation of the phase diagram, surface tension and engaged binding sites per protein through a local order parameter, are provided in the Supporting Information document.

## Acknowledgements

This project has received funding from the European Research Council (ERC) under the European Union Horizon 2020 research and innovation programme (grant agreement No 803326). J. R. E. acknowledges funding from the Oppenheimer Fellowship and from Emmanuel College Roger Ekins Research Fellowship. I. S. B. acknowledges funding from the Oppenheimer Fellowship, EPRSC Doctoral Programme Training number EP/T517847/1 and Derek Brewer Emmanuel College scholarship. J. A. J. is a Junior Research Fellow at Kings College. This work has been performed using resources provided by the Cambridge Tier-2 system operated by the University of Cambridge Research Computing Service (http://www.hpc.cam.ac.uk) funded by EPSRC Tier-2 capital grant EP/P020259/1.

## Author contributions statement

I. S. B. and J.R.E conceived the project. I. S. B. conducted the simulations. I. S. B. and J.R.E analyzed the results. J.A.J. and R.C. G. contextualized the results. I. S. B. and J.R.E wrote the initial version of the manuscript. All the authors reviewed and edited the manuscript.

