## Supporting Information for "Size conservation emerges spontaneously in biomolecular condensates formed by scaffolds and surfactant clients"

### I. COMPUTING PHASE DIAGRAMS AND SURFACE TENSION

To calculate the coexisting densities of the phase diagrams, we employ the Direct Coexistence method [1–3]. Within this scheme, the two coexisting phases are simulated by preparing periodically extended slabs of the two phases, the condensed and the diluted one, in the same simulation box. We use an implicit solvent model; accordingly, the diluted phase (protein-poor liquid phase) and the condensed phase (protein-rich liquid phase) are effectively a vapour and a liquid phase, respectively. We prepare the initial configurations in the following way: First, we build a cubic box configuration containing all the required proteins (patchy particles) for a given mixture. Then, we run a $NpT$ simulation so that our system condenses. For that purpose, conditions of temperature of $T^* = 0.09$ and pressure $p^* = 0.16$ are enough. We then elongate the box in one direction (say, $x$) by performing an $Np_xT$ simulation at $T^* = 0.09$ and $p^* = -0.16$ until we obtain the desired simulation box geometry. Once DC simulations have reached equilibrium, we compute the density profile along the long axis of the box, and thus, we extract the density of the two coexisting phases (as shown in the Supp. Info of Ref. [4]). From the plateau of the condensed phase and the diluted one, we measure the density (avoiding the interfaces between both phases). To estimate the critical point of the phase diagrams, we use the universal scaling law of coexistence densities near a critical point [5], and the law of rectilinear diameters [6]:

$$\left(\rho_l^*(T^*) - \rho_v^*(T^*)\right)^{3.06} = d\left(1 - \frac{T*}{T_c^*}\right) \tag{1}$$

$$(\rho_l^* + \rho_v^*)/2 = \rho_c^* + s_2(T_c^* - T*) \tag{2}$$

where $\rho_l^*$, $\rho_v^*$ refer to the reduced coexisting densities of the condensed and diluted phases respectively, while $\rho_c^*$ is the critical reduced density, $T_c^*$ is the reduced critical temperature, and $d$ and $s_2$ are fitting parameters.

From Direct Coexistence simulations, we can also obtain the surface tension of the condensates by using the following expression:

$$\gamma = \int_{-\infty}^{\infty} [p_N(x) - p_T(x)]\, dx = \frac{L_x}{2N}(\overline{p}_N - \overline{p}_T) \tag{3}$$

where $L_x$ is the long side of the simulation box (perpendicular to the slab interfaces in our simulations), N is the number of droplets in our system, and $p_N$ and $p_T$ are the normal an tangential components of the pressure tensor with respect to the interfaces of the system (note that the tangential component must be averaged over the two tangential directions). In Figure 1 of the main text, we also fit our data of $\gamma$ assuming the following scaling with temperature $\gamma \propto (T_c^* - T^*)^{1.26}$ [5] as described in the caption to the figure.

### II. COMPUTING THE NUMBER OF ENGAGED BONDS PER PROTEIN AND DENSITY PROFILES

In Figure 4 of the main text, we show two profiles, a scaffold/surfactant density profile (a) and a number of engaged bond per protein (scaffold/surfactant) profile (b), both averaged from the center of mass of the condensate. In order to compute these profiles, we use the same order parameter based on the number of neighbours that we used in our previous work (See Appendix of Ref. [7]). For a cut-off distance of $1.20\sigma$, any protein with at least two neighbours is considered part of the condensed phase. Those proteins that exhibit a condensed-phase-like environment, and that additionally are closer than a distance of $1.20\sigma$, are considered to be part of the same droplet. Afterwards, making use of an algorithm to compute the center of mass of the condensate through the periodic boundary conditions [8], we average the density profile (using both directions) from the slab center of mass along the long axis of our simulation box. To compute the number of engaged bonds per protein profile, we evaluate the number of engaged binding sites per protein as a function of distance from the condensate center of mass. The number of used binding sites per protein is the number of binding sites per molecule with another binding site at a closer distance than $0.145\sigma$. For both measurements, the results are averaged among hundreds of independent configurations once the system has reached equilibrium.


[1] A. J. Ladd and L. V. Woodcock, Chemical Physics Letters **51**, 155 (1977).
[2] R. García Fernández, J. L. F. Abascal, and C. Vega, The Journal of Chemical Physics **124**, 144506 (2006).
[3] J. R. Espinosa, E. Sanz, C. Valeriani, and C. Vega, Journal of Chemical Physics **139** (2013), 10.1063/1.4823499.
[4] J. A. Joseph, J. R. Espinosa, I. Sanchez-Burgos, A. Garaizar, D. Frenkel, and R. Collepardo-Guevara, Biophysical Journal **120**, 1219 (2021).
[5] J. S. Rowlinson and B. Widom, *Molecular theory of capillarity* (Courier Corporation, 2013).
[6] J. A. Zollweg and G. W. Mulholland, The Journal of Chemical Physics **57**, 1021 (1972).
[7] I. Sanchez-Burgos, J. R. Espinosa, J. A. Joseph, and R. Collepardo-Guevara, Biomolecules **11**, 278 (2021).
[8] L. Bai and D. Breen, Journal of Graphics Tools **13**, 53 (2008).
